# Taxonomic sampling and rare genomic changes overcome long-branch attraction in the phylogenetic placement of pseudoscorpions

**DOI:** 10.1101/2020.11.18.389098

**Authors:** Andrew Z. Ontano, Guilherme Gainett, Shlomi Aharon, Jesús A. Ballesteros, Ligia R. Benavides, Kevin F. Corbett, Efrat Gavish-Regev, Mark S. Harvey, Scott Monsma, Carlos E. Santibáñez-López, Emily V.W. Setton, Jakob T. Zehms, Jeanne A. Zeh, David W. Zeh, Prashant P. Sharma

**Affiliations:** Department of Integrative Biology, University of Wisconsin-Madison, Madison, WI, USA 53706; National Natural History Collections, The Hebrew University of Jerusalem, Jerusalem, Israel 9190401; Department of Organismic and Evolutionary Biology, Harvard University, Cambridge, MA, USA 02138; Collections & Research, Western Australian Museum, 49 Kew Street, Welshpool, Western Australia 6106, Australia; Lucigen Corporation, 2905 Parmenter Street, Middleton, WU, USA 53562.; Department of Biology, Eastern Connecticut State University, 83 Windham Street, Willimantic, CT 06226, USA; Department of Biology and Program in Ecology, Evolution & Conservation Biology, University of Nevada, Reno, NV 89557, USA

**Keywords:** arachnids, ohnologs, supermatrix, species tree reconciliation, microRNA

## Abstract

Long-branch attraction is a systematic artifact that results in erroneous groupings of fast-evolving taxa. The combination of short, deep internodes in tandem with LBA artifacts has produced empirically intractable parts of the Tree of Life. One such group is the arthropod subphylum Chelicerata, whose backbone phylogeny has remained unstable despite improvements in phylogenetic methods and genome-scale datasets. Pseudoscorpion placement is particularly variable across datasets and analytical frameworks, with this group either clustering with other long-branch orders or with Arachnopulmonata (scorpions and tetrapulmonates). To surmount LBA, we investigated the effect of taxonomic sampling via sequential deletion of basally branching pseudoscorpion superfamilies, as well as varying gene occupancy thresholds in supermatrices. We show that concatenated supermatrices and coalescent-based summary species tree approaches support a sister group relationship of pseudoscorpions and scorpions, when more of the basally branching taxa are sampled. Matrix completeness had demonstrably less influence on tree topology. As an external arbiter of phylogenetic placement, we leveraged the recent discovery of an ancient genome duplication in the common ancestor of Arachnopulmonata as a litmus test for competing hypotheses of pseudoscorpion relationships. We generated a high-quality developmental transcriptome and the first genome for pseudoscorpions to assess the incidence of arachnopulmonate-specific duplications (e.g., homeobox genes and miRNAs). Our results support the inclusion of pseudoscorpions in Arachnopulmonata, as the sister group of scorpions. Panscorpiones (**new name**) is proposed for the clade uniting Scorpiones and Pseudoscorpiones.

## Introduction

The advent of current generation sequencing technologies has greatly benefitted the practice of molecular systematics. However, certain recalcitrant nodes in the Tree of Life remain staunchly unresolved despite the quantity of sequence data deployed to address phylogenetic relationships. Among the most intractable empirical problems in phylogenetics are nodes characterized by the combination of (1) ancient and rapid diversification, and (2) accelerated evolution of multiple ingroup lineages, exacerbating long branch attraction artifacts (Bergsten 2005; Rokas and Carroll 2006; King and Rokas 2017). The combination of these characteristics is difficult to overcome even with genome-scale datasets, due to homoplasy accrued over millions of years of evolutionary history, conflicting evolutionary signals in data partitions, systematic bias, and the lack of external arbiters to evaluate appropriateness of substitution and rate heterogeneity models. Within animals, examples of such problematic nodes include the base of Metazoa, Bilateria, the superclades Lophotrochozoa and Ecdysozoa, and internal relationships of many diverse phyla (Borner et al. 2014; Kocot et al. 2016; Feuda et al. 2017; Simion et al. 2017; Marlétaz et al. 2019; Laumer et al. 2019).

The basal phylogeny of the arthropod subphylum Chelicerata remains particularly recalcitrant to resolution despite the application of genome-scale phylogenomic datasets (Sharma et al. 2014a; Ballesteros and Sharma 2019; Ballesteros et al. 2019; Lozano-Fernández et al. 2019). The basal diversification of this group and the crown age of many orders dates to the early Paleozoic (Lozano-Fernández et al. 2020). Within chelicerates, at least three orders exhibit the characteristics of long-branch orders (Acariformes, Parasitiformes, Pseudoscorpiones), with Solifugae and Palpigradi also prone to unstable placement, as inferred from taxon deletion experiments and assessments of topological stability (Sharma et al. 2014a; Ballesteros and Sharma 2019; Ballesteros et al. 2019). Moreover, extinction has asymmetrically affected different branches in the chelicerate tree, resulting in both relictual orders as horseshoe crabs, as well as several extinct orders. As a result, basic questions about the evolutionary history of Chelicerata remain controversial, namely, the monophyly of Arachnida (the terrestrial chelicerates; Ballesteros and Sharma 2019; Ballesteros et al. 2019; Lozano-Fernández et al. 2019). Even in datasets that support arachnid monophyly, relationships between chelicerate orders are highly unstable from one dataset to the next, with the exception of the basal split between Pycnogonida (sea spiders) and the remaining chelicerates, Tetrapulmonata (a group of arachnid orders that bear four book lungs; Pepato et al. 2010; Regier et al. 2010; Sharma et al. 2014a; Ballesteros and Sharma 2019; Ballesteros et al. 2019; Lozano-Fernández et al. 2019; Howard et al. 2020), and the robust recovery of Arachnopulmonata (Scorpiones + Tetrapulmonata; Sharma et al. 2014a; Ballesteros and Sharma 2019; Ballesteros et al. 2019; Lozano-Fernández et al. 2019; Howard et al. 2020).

In addition to these phylogenomic analyses, Arachnopulmonata is also supported by analyses of genome architecture, as both spiders and scorpions share a partial or whole genome duplication. This inference is evidenced by retention of duplicated copies of numerous developmental patterning genes and microRNAs, to the exclusion of groups like Opiliones (harvestmen) and Acari (Schwager et al. 2007; Sharma et al. 2014b; Sharma et al. 2016; Leite et al. 2016; Schwager et al. 2017; Leite et al. 2018). Moreover, exploratory analyses of gene trees and embryonic gene expression patterns in spiders, scorpions, and harvestmen have shown that the duplicated copies of arachnopulmonate leg-patterning genes also retain expression domains that reflect the evolutionary history of shared whole genome duplication (WGD; Nolan et al. 2020; Gainett and Sharma 2020). The systemic duplication of developmental patterning genes and gene expression patterns together constitute a highly complex character that unites Arachnopulmonata (Leite et al. 2016; Nolan et al. 2020; Gainett and Sharma 2020; Gainett et al. 2020), but the putative incidence of this phenomenon has not been assessed in many chelicerate orders, most of which lack genomic and functional genetic resources (Garb et al. 2018).

One potential solution to overcoming long-branch attraction includes the expansion of taxonomic sampling, which serves to “break” long branches and improve the estimation of parameters of substitution models. While recent efforts have targeted improving taxonomic representation of the acarine orders in phylogenetic datasets (Acariformes and Parasitiformes; Arribas et al. 2019; Charrier et al. 2019), only recently has phylogenomic sampling of Pseudoscorpiones successfully sampled all major extant lineages (Benavides et al. 2019). Intriguingly, in phylogenetic studies that have broadly sampled pseudoscorpions and scorpions, pseudoscorpions are frequently recovered as either sister group to Arachnopulmonata (Sharma et al. 2015a) or as sister group to scorpions (Sharma et al. 2018; Benavides et al. 2019), although these works lacked complete representation of all chelicerate orders. In works assessing chelicerate phylogeny broadly, pseudoscorpion placement has proven unstable or unsupported, either clustering with the Acari or with arachnopulmonates (Sharma et al. 2014a; Arribas et al. 2019; Ballesteros and Sharma 2019; Ballesteros et al. 2019; Lozano-Fernández et al. 2019). In these works, taxonomic representation of Pseudoscorpiones has nevertheless been limited, often to a subset of derived lineages.

To assess these competing hypotheses for pseudoscorpion placement in the chelicerate tree of life, we established a phylogenomic dataset of Chelicerata broadly sampling all major lineages of Pseudoscorpiones. We simulated the effect of incomplete taxonomic sampling by sequentially pruning basally branching lineages of pseudoscorpions and assessed the effect on the inferred tree topology using different analytical approaches to phylogenetic reconstruction. Furthermore, we reasoned that if Pseudoscorpiones is nested within Arachnopulmonata, then it follows they should share the systemic duplications of developmental patterning genes previously demonstrated for scorpions and spiders (Leite et al. 2016, 2018). Here we show that expanded taxonomic sampling of pseudoscorpions, systemic homeobox gene duplications, tree topologies of benchmarked ohnologs of developmental patterning genes, and duplications of miRNAs, all support the hypothesis that pseudoscorpions are nested within Arachnopulmonata as the sister group of scorpions.

## Results

### Phylogenomic results

To assess matrix completeness and denser taxonomic sampling as explanatory processes for the unstable phylogenetic placement of pseudoscorpions, we assembled a dataset of 132 Panarthropoda, including 40 pseudoscorpion libraries previously generated by Benavides et al. (2019), which represent all pseudoscorpion superfamilies (fig. 2a; table S1). Orthologs analyzed in this study consisted of the Benchmarked Universal Single Copy Orthologs of Arthropoda (BUSCO-Ar) discovered in the taxa surveyed. We constructed six matrices ranging in gene occupancy thresholds of 80% (248 BUSCO loci) to 55% (1002 BUSCO loci); we denote these as G_1_ to G_6_, in order of increasing matrix length (Fig. 2b). For each of these six matrices, we additionally pruned basally branches lineages within Pseudoscorpiones, with reference to Cheliferoidea. This superfamily was selected as the distal-most taxon because it was represented by the most exemplars of any pseudoscorpion superfamily (12 transcriptomes), ensuring that the order would be well represented across all super matrices, despite the pruning of other lineages (table S2). For each matrix, we performed maximum likelihood (ML) searches and assessed phylogenetic placement of pseudoscorpions as sister group to scorpions, sister group to Arachnopulmonata (*sensu* Sharma et al. 2014a) or sister group to one or both of the long-branch acarine orders (Acariformes and Parasitiformes). Six branches were sequentially pruned; we denote these datasets as *T*_*−1*_ to *T*_*−6*_, in order of increasing branch pruning (Fig. 2a, 2b)

**Figure 1.**
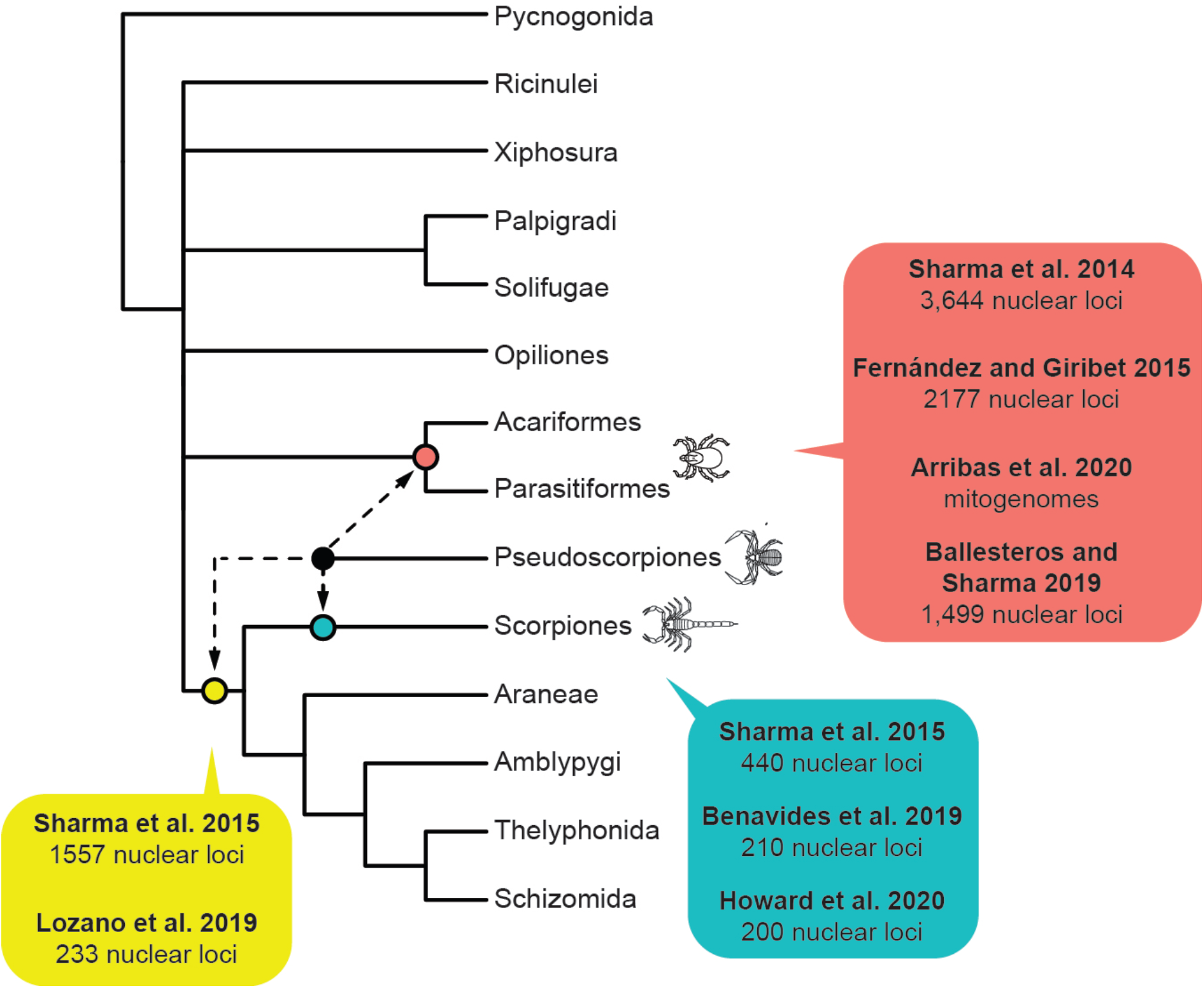
Summary tree topology of Chelicerata showing relationships of orders. Phylogeny based on Ballesteros et al. (2019). Dotted lines for Pseudoscorpiones show alternative placements of this order in selected historical phylogenetic analyses.

**Figure 2.**
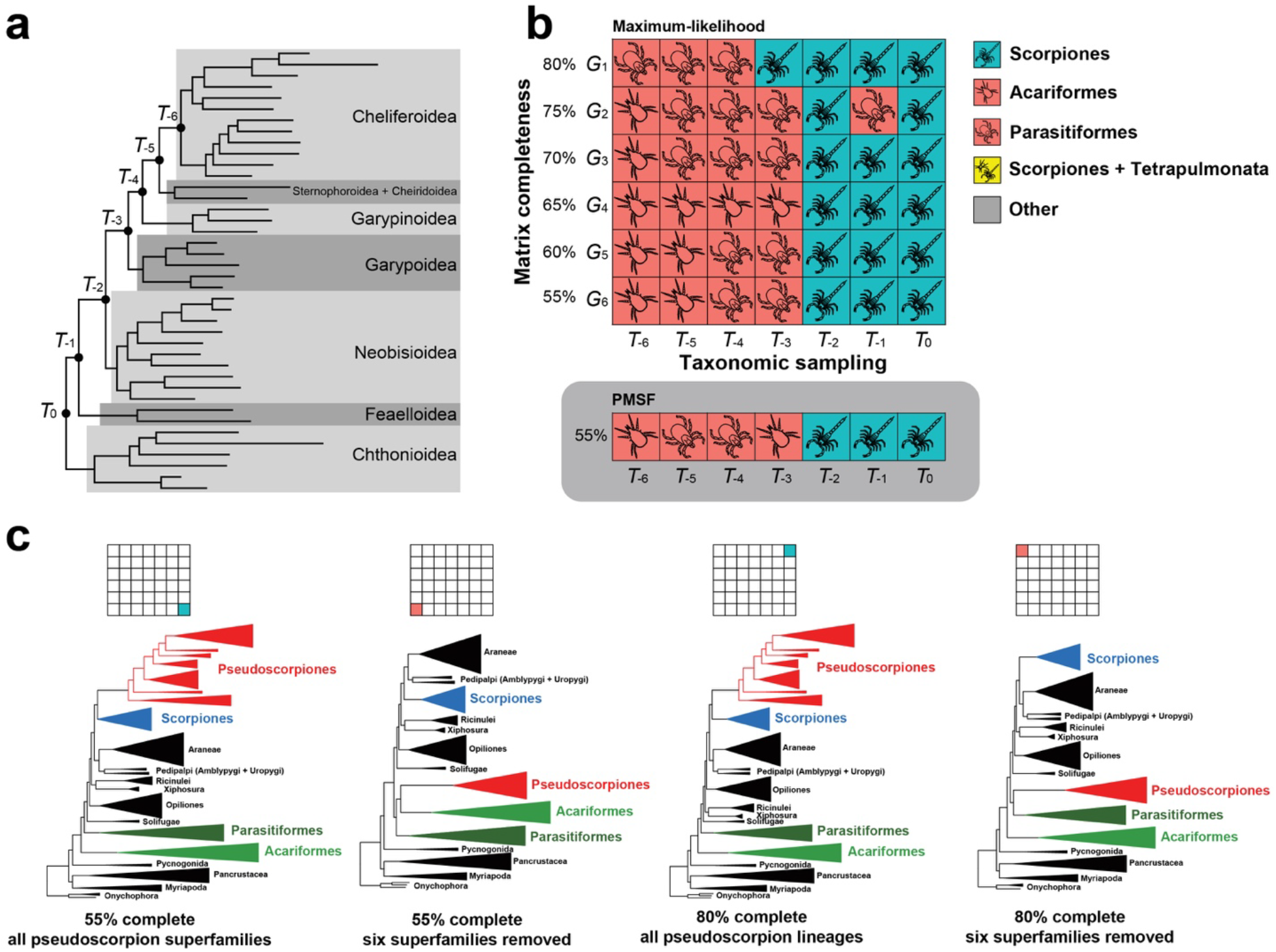
Depth of taxonomic sampling is more influential than matrix completeness in supermatrix analyses of pseudoscorpion placement. (a) Internal phylogeny of Pseudoscorpiones showing major taxonomic groups. Notations on nodes indicate taxon subsets obtained by sequential pruning of branches. (b) Sensitivity plot of 42 phylogenomic matrices assembled by varying gene occupancy (y-axis) and taxonomic sampling (x-axis). Colors of squares correspond to the sister group of Pseudoscorpiones obtained in each maximum likelihood analysis. Gray background: Analysis of largest matrices (1002 loci) under PMSF model. (c) Selected tree topologies showing the dynamics of pseudoscorpion instability as a function of taxonomic sampling.

Matrices retaining all superfamilies of pseudoscorpions (i.e., unpruned datasets) consistently recovered the relationship Pseudoscorpiones + Scorpiones, regardless of matrix completeness. ML tree topologies of pruned taxon subsets *T*_*−1*_ and *T*_*−2*_ similarly recovered the relationship Pseudoscorpiones + Scorpiones, excepting matrix *G*_*2*_ • *T*_*−1*_, which recovered an unsupported relationship of Pseudoscorpiones + Parasitiformes (ultrafast bootstrap resampling frequency [BS] = 43%).

Inversely, matrices exhibiting pruning of the three most basally branching pseudoscorpion lineages (Chthonioidea, Feaelloidea, and Neobisioidea) recovered ML tree topologies wherein pseudoscorpions were sister group to either Parasitiformes or Acariformes, regardless of matrix completeness (*T*_*−3*_ matrices). Further pruning of basally branching pseudoscorpions generally also incurred this tree topology (*T*_*−4*_ to *T*_*−6*_ matrices), with the exception of matrix *G*_*1*_ • *T*_*−3*_ (fig. 2b). No matrix recovered the relationship of pseudoscorpions as sister group to Arachnopulmonata (*sensu* Sharma et al. 2014a).

Partitioned model ML analyses have sometimes been criticized less inaccurate than site heterogeneous models, although these inferences have often been grounded in preconceptions of true phylogenetic relationships based on traditional phylogenetic hypotheses (e.g., Wang et al. 2019). Simulations have previously shown that CAT+GTR and partitioned ML analyses are comparably accurate, with both of these outperforming CAT-F81 (sometimes referred to as CAT-Poisson) with respect to topological accuracy (Halanych et al. 2017). However, CAT+GTR models are notoriously difficult to implement in a Bayesian framework, due to excessive computational times for real datasets (i.e., > 100 taxa, > 500 genes), and numerous published analyses using PhyloBayes-mpi have exhibited failure to converge (defined as ESS > 200; *maxdiff* < 0.01), especially for chelicerate phylogeny (Sharma et al. 2014a; Ballesteros and Sharma 2019; Ballesteros et al. 2019; Lozano-Fernández et al. 2019; Howard et al. 2020). As a workaround, we assessed the performance of the posterior mean site frequency (PMSF) model (LG + C20 + F + Γ), a mixture model alternative to the CAT implementation. This model was trialed for the *G*_*6*_ family of matrices, which constitute the largest matrices we analyzed (1002 genes). The pattern of tree topologies recovered reflected the same outcome as the partitioned model analyses, with *T*_*−3*_ to *T*_*−6*_ matrices recovering Pseudoscorpiones as clustering with one of the acarine orders, and *T*_*0*_ to *T*_*−2*_ matrices recovered Pseudoscorpiones + Scorpiones (Fig. 2b).

Relationships among other chelicerate taxa largely reflected the outcomes of our previous works (Ballesteros and Sharma 2019; Ballesteros et al. 2019) and are not discussed in detail here. Notably, we never recovered the monophyly of Acari or Arachnida; across all analyses, Xiphosura was recovered as the sister group of Ricinulei (70/91 analyses), or to Ricinulei + Solifugae (21/91 analyses), as previously reported (Ballesteros and Sharma 2019; Ballesteros et al. 2019).

### Nodal support

Ultrafast bootstrap resampling frequencies were used to estimate support for competing hypotheses for the phylogenetic placement of Pseudoscorpiones, across the 42 matrices analyzed (Fig. 3a). Across all levels of matrix completeness, support for Pseudoscorpiones + Scorpiones was negligible (< 10%) for *T*_*−3*_ to *T*_*−6*_ matrices, but increased dramatically upon including Neobisioidea (*T*_*−2*_ matrices). Increase in nodal support for Pseudoscorpiones + Scorpiones was not monotonic, as sampling of Feaelloidea and Chthonioidea resulted in some variability in bootstrap frequency (Fig. 3a). The nodal support trajectories were identical for the hypotheses Pseudoscorpiones + Scorpiones and Pseudoscorpiones + Arachnopulmonata. This result reflects in part the nestedness of the two hypotheses (i.e., Scorpiones is nested within Arachnopulmonata).

**Figure 3.**
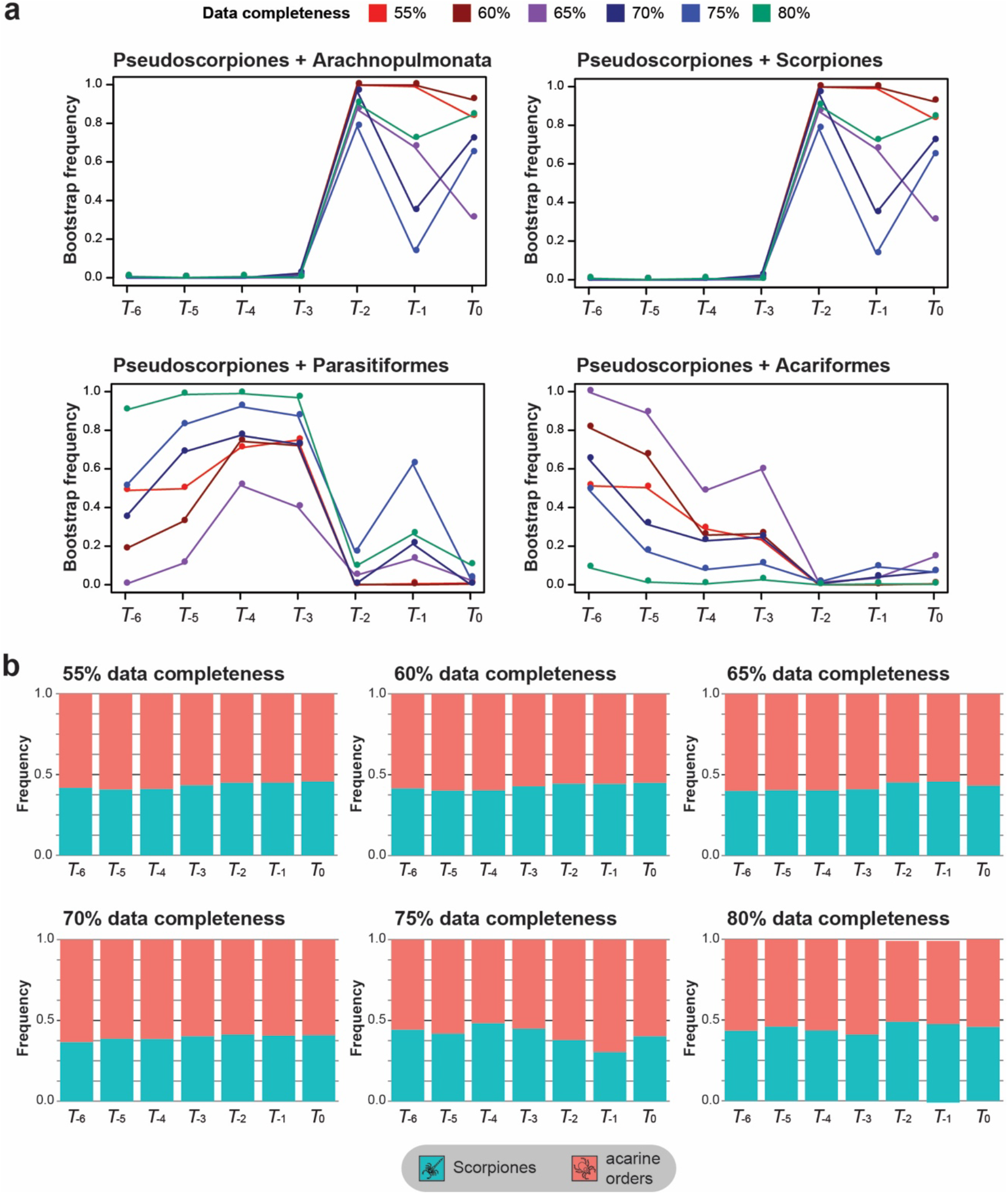
Depth of taxonomic sampling affects supermatrix nodal support, but not per-locus support. (a) Nodal support frequency for competing hypotheses of pseudoscorpion placement as a function of taxonomic sampling and matrix completeness. (b) Proportion of loci favoring Pseudoscorpiones + Scorpiones versus Pseudoscorpiones + either acarine order under a ΔGLS framework, as a function of taxonomic sampling and matrix completeness.

By contrast, support for Pseudoscorpiones as the sister group of either Parasitiformes or Acariformes showed the opposite trend, with better representation of basally branching pseudoscorpion groups resulting in lower nodal support for pseudoscorpions clustering with either of these groups. For taxon subsets with the least representation of basally branching pseudoscorpions (*T*_*−4*_ to *T*_*−6*_ matrices), the most complete matrices recovered high support values for Pseudoscorpiones + Parasitiformes, whereas matrices with intermediate gene occupancy thresholds (60-70%, or *G*_*3*_ to *G*_*5*_ matrices) recovered high support values for Pseudoscorpiones + Acariformes.

### Gene trees, ΔGLS, and species tree reconstruction

Approaches to inferring species tree using gene trees have been shown to be powerful predictors of phylogenetic accuracy, but these methods are predicated on the accuracy of the underlying gene tree set. To assess whether improving taxonomic sampling of a long-branch taxon also affects phylogenetic signal at the level of gene trees, we calculated gene-wise log-likelihood scores (ΔGLS) on gene trees corresponding to each of the 42 matrices. ΔGLS assesses the likelihood of each gene given two competing tree topologies, across all genes in a dataset (Shen et al. 2017). We generated ΔGLS distributions for the two competing hypotheses of pseudoscorpion placement (clustering with scorpions versus clustering with either acarine order).

We observed minimal effects of taxon pruning in the largest matrices (G_5_ and G_6_), and no consistent trends in the distribution of genes favoring either competing hypothesis, across the ΔGLS distributions of 42 analyses (fig. 3b). Magnitudes of log likelihood favoring either hypothesis were also not consistently affected (fig. S1). These results suggest that increasing taxonomic sampling of a long-branch lineage does not greatly alter the distribution of phylogenetic signal at the level of individual gene trees.

To assess whether the intransigence of ΔGLS distributions to taxonomic sampling has downstream effects on methods of phylogenetic reconstruction, especially those that use the multispecies coalescent model, we reconstructed species trees from gene trees using ASTRAL v.3 (Zhang et al. 2018). We discovered no clear difference between the performance of ASTRAL v.3 versus concatenation based approaches, with respect to the tree topology recovered as a function of the number of basal branches pruned (fig. 4). Generally, *T*_*0*_ to *T*_*−2*_ matrices recovered the relationship Pseudoscorpiones + Scorpiones, whereas *T*_*−3*_ to *T*_*−6*_ matrices again recovered Pseudoscorpiones as clustering with the acarine orders. The exceptions were matrices *G*_*2*_ • *T*_*−1*_ and *G*_*4*_ • *T*_*0*_, which recovered pseudoscorpions as the sister group of Parasitiformes or in a grade at the base of Chelicerata, respectively.

**Figure 4.**
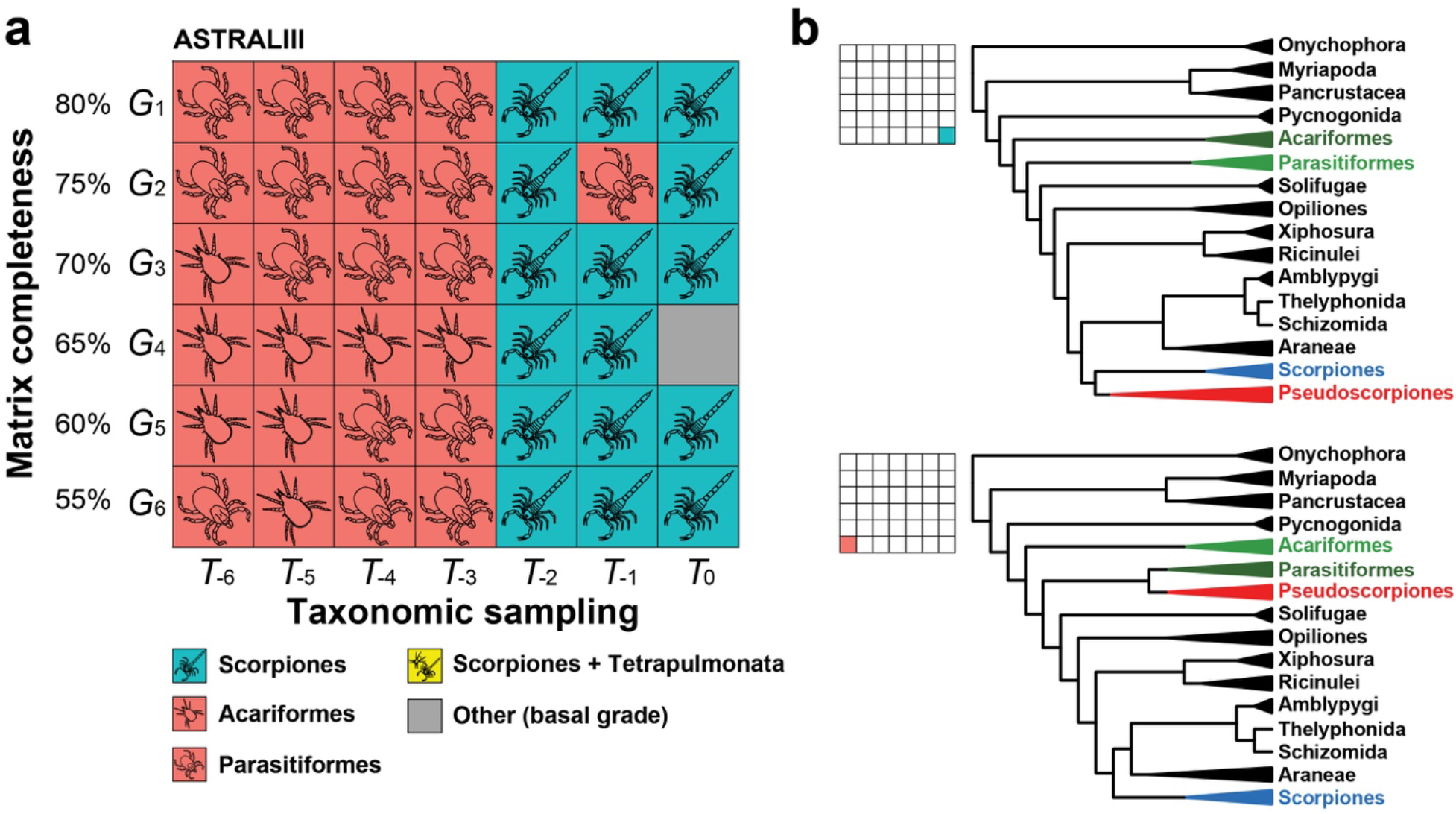
Depth of taxonomic sampling is more influential than matrix completeness in ASTRAL analyses of pseudoscorpion placement. (a) Sensitivity plot of 42 phylogenomic matrices assembled by varying gene occupancy (y-axis) and taxonomic sampling (x-axis). Selected tree topologies showing the dynamics of pseudoscorpion instability as a function of taxonomic sampling.

**Figure 5.**
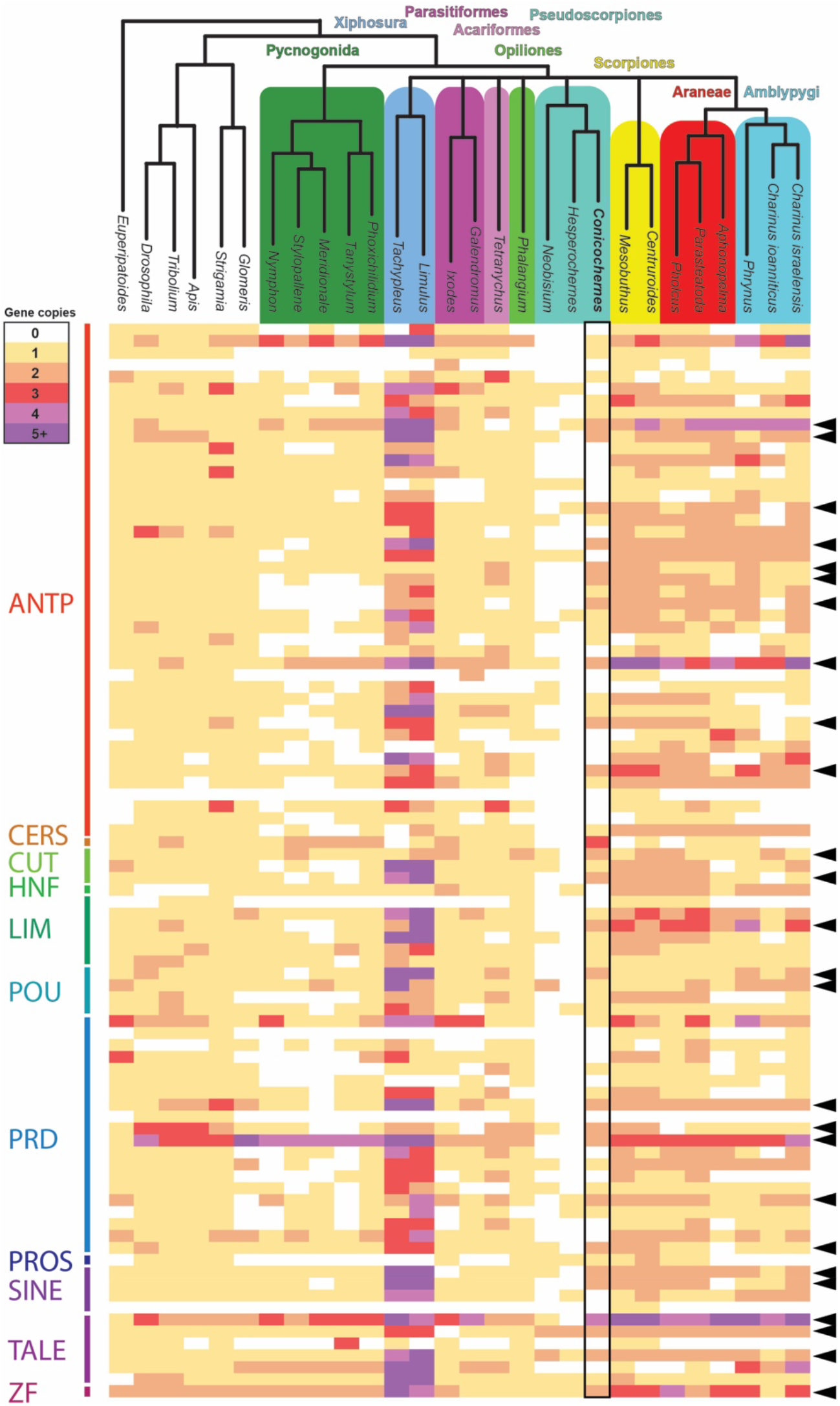
Comparison of homeobox repertoires for 26 panarthropods supports retention of duplications in pseudoscorpions that are shared with arachnopulmonates. Rows correspond to individual homeobox genes. Colors correspond to numbers of paralogs. Black arrows to the right indicate duplications in at least one pseudoscorpion exemplar that is also shared by at least one arachnopulmonate.

**Figure 6.**
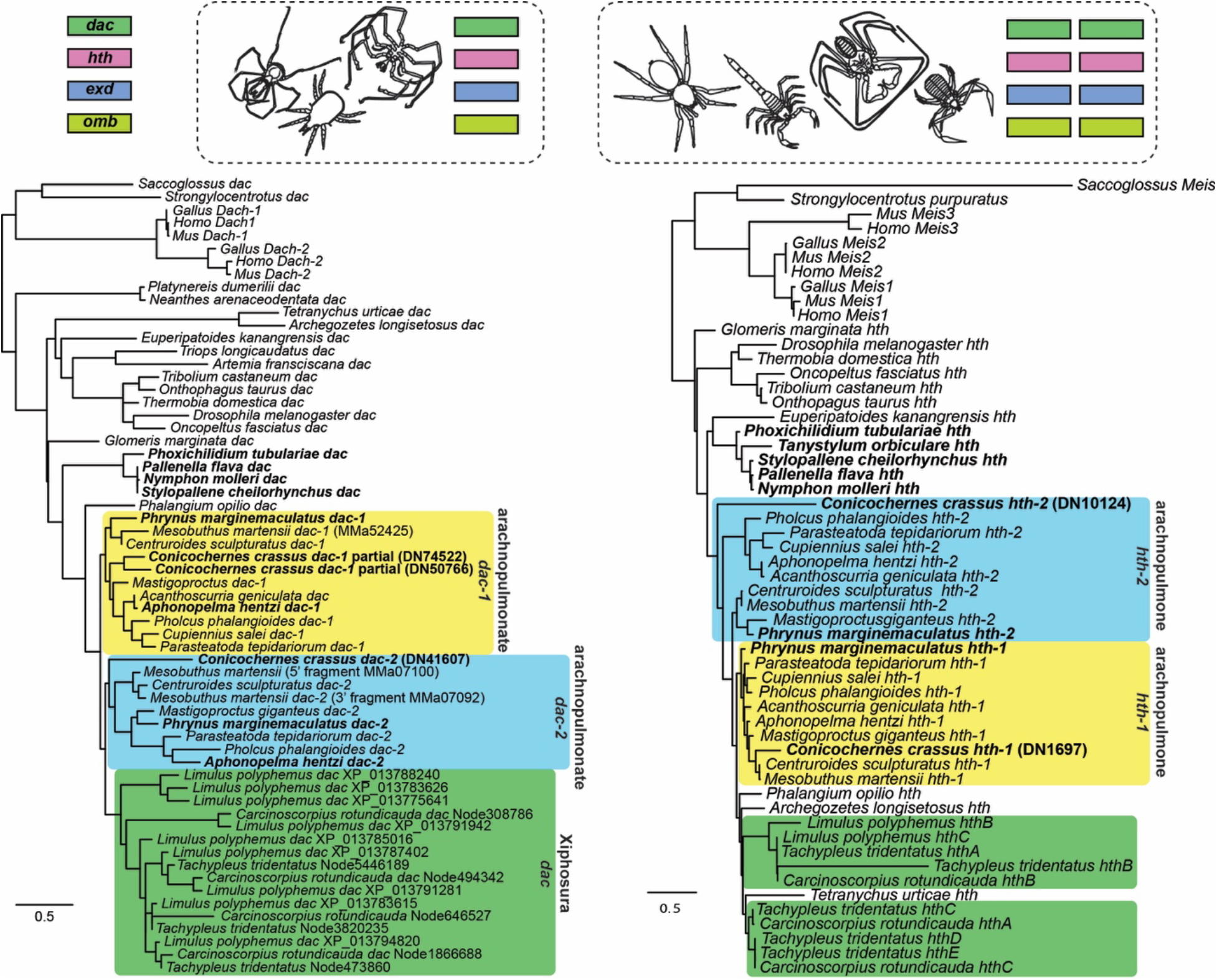
Pseudoscorpions possess two copies of four developmental patterning genes with benchmarked, arachnopulmonate-specific divergence of spatiotemporal expression domains. Above: Single copy orthologs of *dac*, *hth*, *exd*, and *omb* were recovered from genomic resources for sea spiders, harvestmen, and Acariformes, whereas two copies of each gene were recovered for scorpions, tetrapulmonates, and pseudoscorpions. Below left: Maximum likelihood gene tree topology of the medial leg gap gene *dac*. Below right: Maximum likelihood gene tree topology of the proximal leg gap gene *hth*. Note the clustering of pseudoscorpion copies within arachnopulmonate clusters. *dac-1* of *C. crassus* was recovered as two nonoverlapping fragments.

ASTRAL analyses never recovered the monophyly of Arachnida or Acari.

### Duplications of homeobox genes

As an external arbiter of the two competing hypotheses of pseudoscorpion relationships, we generated a developmental transcriptome of the West Australian chernetid *Conicochernes crassus*. Homeobox gene surveys of developmental transcriptomes and/or genomes have previously been shown to be faithful readouts of whole genome duplication in Chelicerata. Whole genome duplications are inferred to have occurred in the common ancestor of Arachnopulmonata (one event) and independently in the Xiphosura (twofold or threefold WGD); groups like mites, ticks, and harvestmen do not exhibit these shared duplications (Sharma et al. 2014b; Kenny et al. 2015; Sharma et al. 2015a; Kenny et al. 2016; Schwager et al. 2017; Leite et al. 2016; Leite et al. 2018; Shingate et al. 2020). A previous comprehensive analysis of homeobox genes by Leite et al. (2018) showed that the retention of duplicates is systemic in two arachnopulmonate lineages (spiders and scorpions), an inference subsequently supported by the first whip spider developmental transcriptomes (Gainett and Sharma 2020; Gainett et al. 2020) and by embryonic gene expression data (Nolan et al. 2020; Gainett and Sharma 2020). However, this survey of homeobox duplications omitted key groups, such as Xiphosura and Parasitiformes (Leite et al. 2018). Curiously, Leite et al. (2018) had indeed sampled two pseudoscorpion species, but recovered few homeobox genes for these taxa, likely owing to the sampling of postembryonic stages rather than embryos; in scorpions, developmental transcriptomes have been shown to recover far more duplicated homeobox genes than adult transcriptomes (Sharma et al. 2014b, 2015a).

We therefore assembled a dataset of 26 Panarthropoda, sampling genomes or developmental transcriptomes of all three major lineages of Arachnopulmonata *sensu* Sharma et al. (2014a) (i.e., spiders, scorpions, and Pedipalpi [Amblypygi + Thelyphonida + Schizomida]), as well as mites, ticks, harvestmen, horseshoe crabs, and sea spiders. This dataset leveraged recent developmental genetic resources generated by us for several non-model chelicerate groups, such as mygalomorph spiders, whip spiders, harvestmen, and sea spiders (Sharma et al. 2012; Gainett and Sharma 2020; Gainett et al. 2020; Ballesteros et al. 2020). We included in our analysis two adult transcriptomes of pseudoscorpions previously analyzed by Leite et al. (2018), which had been shown to harbor few homeobox genes and exhibited short contigs for many homeobox homologs. Outgroup datasets consisted of an onychophoran embryonic transcriptome and genomes of Mandibulata.

In contrast to previous analyses of adult pseudoscorpion transcriptomes (*Hesperochernes* sp. and *Neobisium carcinoides* in Leite et al. 2018), our analysis of the first pseudoscorpion developmental transcriptome recovered homologs of 56 homeobox genes in *C. crassus*. Of these, 26 exhibited duplications in at least one of the three pseudoscorpion exemplars that were also found in at least one scorpion or one tetrapulmonate), with clear evidence of paralogy (i.e., overlapping peptide sequences exceeding 100 amino acids in length that exhibited multiple substitutions between duplicate pairs).

All ten Hox genes ancestral to Panarthropoda are known to be duplicated in scorpions and spiders, with embryonic expression patterns reflecting the shared duplication (Schwager et al. 2007; Sharma et al. 2014b; Schwager et al. 2017). Recent work has shown that the common ancestor of Amblypygi (whip spiders) likely also exhibited two copies of each Hox gene (Gainett and Sharma 2020). However, the previous homeobox survey of adult pseudoscorpion transcriptomes had only recovered five of the ten Hox genes, with none of these duplicated (Leite et al. 2018). By contrast, we discovered eight of the ten Hox homologs in the developmental transcriptome of *C. crassus* (all but *Hox3* and *Sex combs reduced*). Of these eight, five exhibited duplications: *labial*, *Deformed*, *fushi tarazu*, *Antennapedia*, and *abdominal-A*.

Other well-characterized embryonic patterning genes among the homeobox family that were duplicated in both pseudoscorpions and arachnopulmonates included the Six genes (e.g., *sine oculis*; *Optix*; Gainett et al. 2020), central nervous system patterning genes (e.g., *empty spiracles*; *Pax3/7*) appendage patterning genes (e.g., *homothorax*; *extradenticle*; ref. Nolan et al. 2020), and segmentation cascade genes (e.g., *engrailed*; *orthodenticle*). Enumeration of the homeobox homologs across the 26 species is provided in table S3.

By comparison to pseudoscorpions, we did not detect systemic duplications of homeobox genes (i.e., suggestive of shared whole genome duplication with arachnopulmonates) in Acariformes, Parasitiformes, Opiliones, or Pycnogonida. As a key example, among these groups of arachnids, duplicates of only two Hox genes were detected in the genome of the mite *Tetranychus urticae* (with these being tandem duplicates on a single Hox cluster; fig. 4a of Grbic et al. 2011). By contrast, tetrapulmonate exemplars new to this analysis (the mygalomorph *Aphonopelma hentzi*; the three Amblypygi species) exhibited the expected trend of retention of homeobox duplicates. Taken together, this survey of homeobox genes suggests that pseudoscorpions were included in the shared whole genome duplication at the base of Arachnopulmonata.

### Gene tree analysis of benchmarked embryonic patterning genes

Whereas embryonic expression data are abundant for spiders, and principally for the model system *Parasteatoda tepidariorum*, they are comparatively few for non-spider chelicerate groups (e.g., Blackburn et al. 2006; Jager et al. 2006; Grbic et al. 2007; Barnett and Thomas 2013; Sharma et al. 2012; Sharma et al. 2014b; Sharma et al. 2015b; Gainett and Sharma 2020). In a recent comparative work, it was shown that four appendage patterning genes known to be duplicated in spiders and scorpions exhibited shared expression patterns that reflected the history of the species tree (i.e., ohnologs of *P. tepidariorum* and the scorpion *C. sculpturatus* exhibited orthologous expression patterns, by comparison to the expression domains of the single-copy homologs of outgroups like harvestmen, mites, and mandibulates) (Nolan et al. 2020). These four genes (*dachshund*, *homothorax*, *extradenticle*, and *optomotor blind*) constitute benchmarked cases of arachnopulmonate ohnologs that have been validated via gene expression surveys, with additional and recent corroboration of this pattern in two of the four genes in the whip spider *P. marginemaculatus* (Gainett and Sharma 2020).

We therefore investigated whether duplicates of these four genes also occurred in the developmental transcriptome of *C. crassus*. To the surveys previously generated by Nolan et al. (2020), we searched for and added homologs of these genes from developmental transcriptomes of the pseudoscorpion, the whip spider species *Phrynus marginemaculatus* (Gainett and Sharma 2020), five sea spider species (Setton et al. 2018; Ballesteros et al. 2020) and the tarantula *A. hentzi* (Setton et al. 2019). We discovered two copies of all four genes in the developmental transcriptome of the pseudoscorpion, except for *dachshund*, wherein three putative homologs were discovered. However, two of these pseudoscorpion *dachshund* fragments were non-overlapping, suggesting that only two copies of *dachshund* are present in this transcriptome (comparable to the case of *Mesobuthus martensii*; Nolan et al. 2020). Similarly, we discovered two copies of these genes in the new arachnopulmonate datasets (whip spiders and the tarantula). By contrast, only one copy of these four genes was discovered in the sea spiders, as with mites, ticks, and harvestmen.

Gene tree analysis of these four genes had previously shown sufficient signal to resolve monophyletic clusters of arachnopulmonate *dac* and *hth* ohnologs (Nolan et al. 2020). Upon reconstructing these two gene trees after adding the pseudoscorpion, the whip spiders, the tarantula, and the sea spiders, we observed each pseudoscorpion paralog clustering with a species arachnopulmonate ohnolog, rather than with the single copy orthologs of acarine taxa. For *dac*, the arachnopulmonate (including pseudoscorpion) clusters were recovered as monophyletic; as previously reported, the horseshoe crab duplications are unrelated to those of Arachnopulmonata (Nolan et al. 2020; Shingate et al. 2020). For *hth*, one arachnopulmonate (including pseudoscorpion) ohnolog (*hth-1*, the ohnolog reflecting the ancestral expression pattern; Nolan et al. 2020) was recovered as monophyletic, whereas the other (*hth-2*, the copy with the derived expression pattern; Nolan et al. 2020) was resolved as a grade. Gene trees of *extradenticle* and *optomotor blind* showed insufficient phylogenetic signal for testing phylogenetic placement, as previously reported (Nolan et al. 2020). These results corroborate the inference that systemic duplication unites pseudoscorpions with Arachnopulmonata.

### Hox genes and microRNA duplications in the pseudoscorpion genome

Embryonic transcriptomes have proven useful for inference of gene duplications, but are inferentially limited in that absence of gene copies cannot be distinguished as the result of gene loss versus absence of expression in the sequenced tissue. As a separate validation of systemic duplication in Pseudoscorpiones, we sequenced and analyzed the draft genome of the species, *Cordylochernes scorpioides* for Hox gene clusters and miRNAs. Due to the fragmentation of the assembly, we were unable to recover more than one Hox gene per scaffold. Nevertheless, we discovered 18 Hox genes in the *C. scorpioides* genome, corresponding to two ohnologs of all Hox genes except for *Hox3* (fig. 7). Together with the homeobox duplications in *C. crassus*, these results are consistent with a shared genome duplication uniting arachnopulmonates and pseudoscorpions.

**Figure 7.**
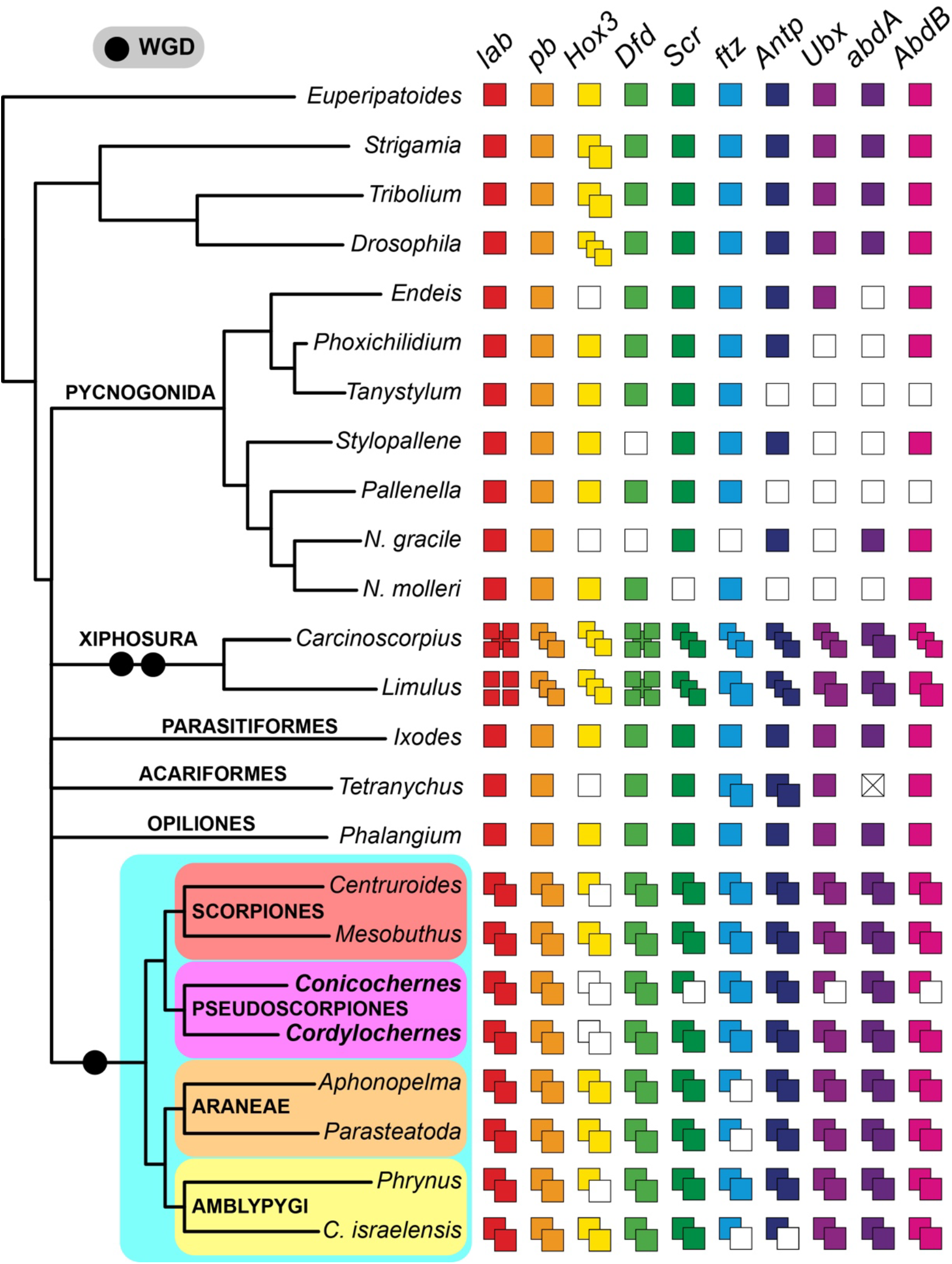
Hox gene complement in the pseudoscorpion genome substantiates evidence of shared whole genome duplication with other Arachnopulmonata. Columns and colored squares correspond to each Hox gene. Unfilled squares correspond to absences, not losses. Cross through *abdA* in *T. urticae* indicates loss of this Hox gene in the mite genome. Note independent twofold whole genome duplication events in Xiphosura (Kenny et al. 2016; Shingate et al. 2020).

MicroRNAs (miRNAs) have been leveraged as rare genomic changes across the metazoan tree of life, with their effectiveness as phylogenetic markers being closely tied to the quality of genomic resources used for miRNA surveys (Tarver et al. 2013; Thomson et al. 2014; Tarver et al. 2018). In Chelicerata, Leite et al. (2016) previously surveyed miRNAs in the genomes of four spiders, a scorpion, a horseshoe crab, five Parasitiformes, and one Acariformes, as well as several outgroup taxa. This survey revealed lineage-specific duplications in *Limulus polyphemus* consistent with twofold whole genome duplication in Xiphosura; duplicated clusters of miRNAs in the spider *P. tepidariorum*, as well as tandem duplications; and a subset of duplicated miRNAs that were shared across spiders and scorpions.

To elucidate if pseudoscorpions exhibit miRNA duplications shared by arachnopulmonates, we expanded the survey of Leite et al. (2016) and searched for miRNAs in the draft genome of the pseudoscorpion, *C. scorpioides* and the genome of the scorpion, *Mesobuthus martensii*. Twenty-six conserved miRNA families were identified in the *C. scorpioides* genome, and another 35 in *M. martensii*. Among them, families iab-4, mir-71, and mir-276 had two or more ortholog copies in Arachnopulmonata, Pseudoscorpiones and Xiphosura (fig. 8). Similarly, we found two members of the families bantam and mir-1 in Scorpiones, Pseudoscorpiones, two spiders and Xiphosura.

**Figure 8.**
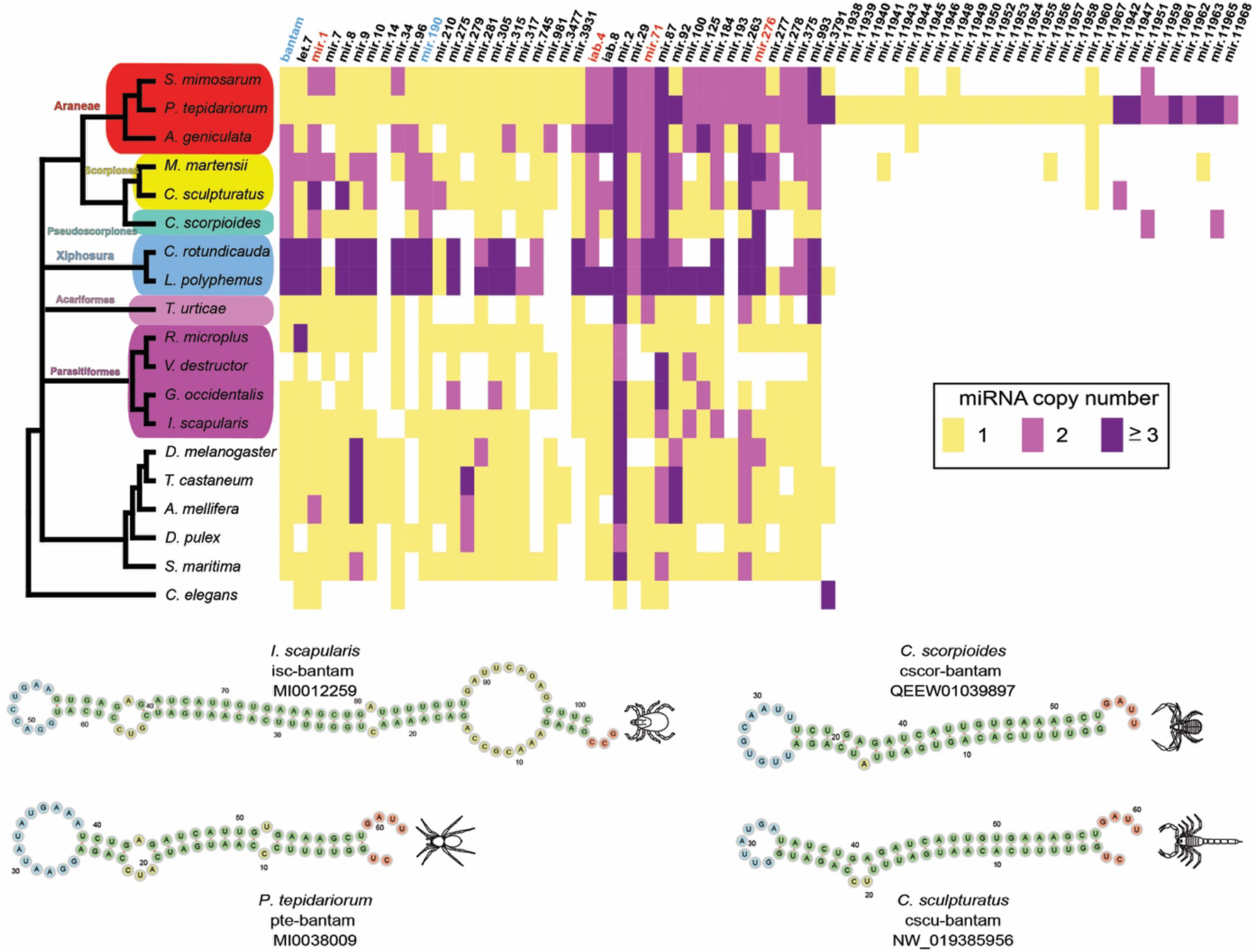
Comparison of miRNA family copy number in *C. scorpioides* and other ecdysozoans supports retention of duplications in pseudoscorpions shared with scorpions and spiders. Columns correspond to individual miRNA families. Colors correspond to numbers of paralogs. miRNA families in red text indicate duplications in scorpions, pseudoscorpions and at least one spider. miRNA families in blue text are duplicated only in scorpions and pseudoscorpions.

Two miRNAs, mir-190 and pte-bantam, were found duplicated only in Scorpiones and *C. scorpioides* (with inferred independent duplications in Xiphosura, fig. 8). Our survey did not recover the presence of miRNA sequences from the families mir-210, mir-275, mir-315, mir-981, mir-277, and mir-11960 (previously reported in genomes of spiders and scorpions). We cannot rule out that these absences are attributable to the incompleteness of the pseudoscorpion genome assembly.

We found no miRNAs unique to Arachnida, nor patterns of duplication consistent with arachnid monophyly.

Taken together, these surveys of miRNA duplication revealed four miRNA duplications supporting the inclusion of pseudoscorpion within arachnopulmonates, and two further duplications supporting the sister relationship of pseudoscorpions and scorpions.

## Discussion

### Consilience of phylogenetic data classes in the placement of pseudoscorpions

Chelicerate higher-level phylogeny is plagued by topological uncertainty, with a subset of orders exhibiting long branch attraction artifacts, as elucidated by taxon deletion experiments (Ballesteros and Sharma 2019; Ballesteros et al. 2019). Barring the monophyly of Euchelicerata (Xiphosura and arachnids), Arachnopulmonata (previously defined as Scorpions + Tetrapulmonata), and relationships within Tetrapulmonata, ordinal relationships in the chelicerate tree of life are highly unstable across phylogenomic datasets. Here, we leveraged the previous discovery of a whole genome duplication subtending the common ancestor of spiders and scorpions to assess competing hypotheses for the placement of pseudoscorpions (Sharma et al. 2014b; Schwager et al. 2017). Taxon-rich analyses of supermatrices as well as reconciliation of gene trees consistently recovered pseudoscorpions as the sister group of scorpions, the hypothesis supported by genome and miRNA duplication. Our taxon deletion experiments reveal that sampling of basally branching lineages in the pseudoscorpion tree of life is key to overcoming long branch attraction artifacts that draw pseudoscorpions together with the acarine orders. Our results are also consistent with the variance of tree topologies in previous chelicerate phylogenetics. Studies that have omitted basally branching pseudoscorpion families, or insufficiently sampled outgroup lineages, recovered Pseudoscorpiones as sister group to, or nested within, Acari (e.g., Sharma et al. 2014a; Arribas et al. 2020). By contrast, phylogenomic works that sampled basal splits within Pseudoscorpiones have recovered support for their placement within Arachnopulmonata (e.g., Benavides et al. 2019; Howard et al. 2020). Our analyses further demonstrate that taxonomic sampling outweighs matrix completeness and analytical approach (supermatrix versus gene tree reconciliation approaches) in achieving phylogenetic accuracy when long branch attraction is incident.

To date, no morphological data matrix has ever recovered the monophyly of Arachnopulmonata (with or without pseudoscorpions), with both older and recent morphological cladistic studies continuing to recover the archaic grouping of Lipoctena (scorpions as the sister group to the remaining arachnid orders; Legg et al. 2013; Lamsdell 2016; Bicknell et al. 2019; Aria and Caron 2019; reviewed by Nolan et al. 2020). Shultz (1990, 2007) presented the first compelling cladistic analyses demonstrating that scorpions are derived within the arachnid tree, a result reflected in another body of recent paleontological investigations (e.g., Garwood and Dunlop 2014; Wang et al. 2018; Huang et al. 2018). In such works, pseudoscorpions have typically been recovered as the sister group of Solifugae (as the clade Haplocnemata), another order exhibiting topological instability (Ballesteros et al. 2019; but see Dunlop et al. 2012). Nevertheless, a sister group relationship of scorpions and pseudoscorpions has previously been tenuously supported by some morphological analyses, namely, the cladistic analysis of Garwood and Dunlop (2014). Subsequent expansion and reuse of this matrix also recovered this relationship (Wang et al. 2018; Huang et al. 2018). However, the recovery of the clade Pseudoscorpiones + Scorpiones as a sister group of Opiliones in those studies is refuted by phylogenomic analyses, developmental gene expression, and genomic architecture (Sharma et al. 2014; Ballesteros et al. 2019; Lozano-Fernandez et al. 2019; Nolan et al. 2020). We therefore observe only partial concordance between our analyses and inferences based on morphological matrices, and only with respect to the placement of pseudoscorpions.

By contrast to morphology, we identified clear and systemic evidence for a shared whole genome duplication in the first developmental transcriptome and genome of two pseudoscorpion exemplars, which is concordant with the hypothesis that pseudoscorpions are derived arachnopulmonates. Surveys of homeobox gene duplication, gene tree topologies of benchmarked arachnopulmonate-specific ohnologs with known spatiotemporal subdivision of embryonic expression domains, and patterns of miRNA duplication all support the inclusion of Pseudoscorpiones within arachnopulmonates, with further evidence from two miRNA families for the clade Pseudoscorpiones + Scorpiones, a clade we term **Panscorpiones**. Henceforth, we redefine Arachnopulmonata to include Pseudoscorpiones.

Due the unanticipated large size of the *C. scorpioides* genome (3.6 Gb), and the ensuing fragmentation of the assembly, we were not able to assess the number of Hox clusters in Pseudoscorpiones, which would constitute an independent test of the hypothesized shared whole genome duplication (but see Hoy et al. 2016 for a case of atomized Hox clusters in a mite). A forthcoming long-read, proximity ligation-based genome assembly of this species is anticipated to inform the ancestral architecture of arachnopulmonate genomes. One additional line of evidence that would support this phylogenetic inference would be embryonic gene expression patterns of ohnologs known to exhibit shared spatiotemporal dynamics in developing appendages of spiders and scorpions (e.g., *dac*; *hth*; Nolan et al. 2020). More recently, evidence from whip spiders (Amblypygi) has additionally supported the inference of conserved expression domains of ohnologs that correspond to gene tree topologies (Gainett and Sharma 2020). While we endeavored to generate expression data for the two copies of the appendage patterning transcription factors *dac*, *hth*, *exd*, and *omb* in *C. crassus*, we encountered technical challenges incurred by cuticle deposition early in pseudoscorpion development, as well as paucity of embryonic tissue. Whole mount in situ hybridization in pseudoscorpion embryos likely requires modified in situ hybridization protocols previously developed for highly sclerotized chelicerate embryos (e.g., sea spiders; Jager et al. 2006). Future efforts must establish a reliable pseudoscorpion model system for testing the downstream hypothesis that expression patterns of pseudoscorpion ohnolog pairs reflect arachnopulmonate-specific patterns. The establishment of a reliable pseudoscorpion model system would constitute a useful comparative data point for assessing the decay of ohnologs’ expression patterns as a function of phylogenetic distance.

### Ancient origins of courtship behavior and brood care in Arachnopulmonata

The recovery of Pseudoscorpiones as the sister group of scorpions markedly alters the reconstruction of several key character in the chelicerate tree of life (fig. 9). Regarding their respiratory system, Pseudoscorpions are reconstructed as arachnopulmonates that have secondarily lost their book lungs; instead, pseudoscorpions typically exhibit two pairs of tracheal tubules opening as spiracles on the third and fourth opisthosomal segments. The evolutionary transition of book lungs to tracheal tubules is broadly associated with miniaturization in other arachnopulmonate orders (Dunlop 2019). For example, in derived spiders, the posterior pair of book lungs is replaced by openings of the tracheal tubules as well, which in turn have a complex evolutionary history within this order (Ramírez et al. 2020). In Schizomida, the posterior pair of respiratory organs is lost altogether (Hansen and Sørensen 1905; Shultz 1990).

**Figure 9.**
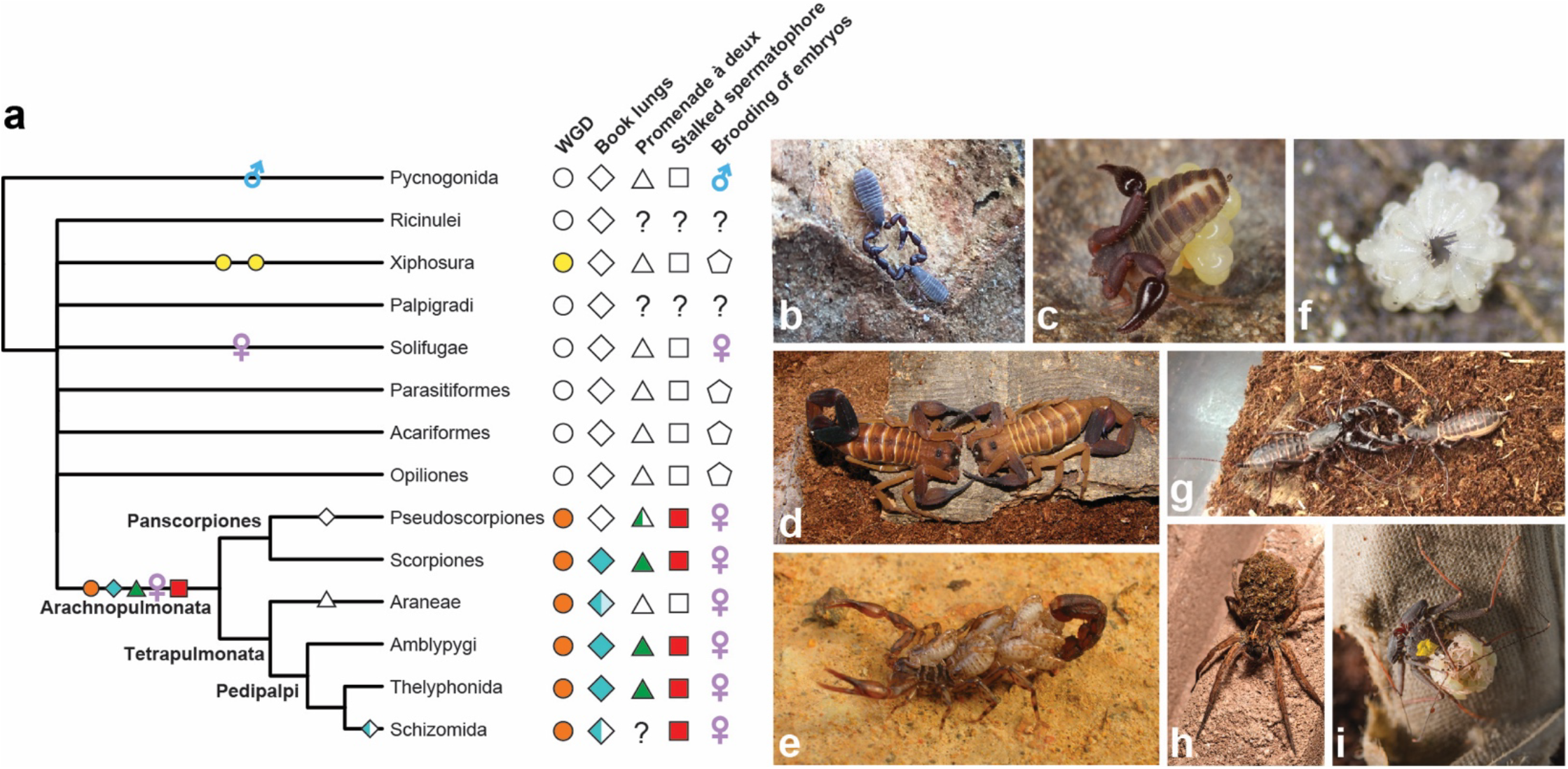
Ancestral state reconstruction of shared genome duplication events and reproductive behaviors in Chelicerata, under accelerated transformation. (a) The revised placement of pseudoscorpions supports a shared origin of courtship behavior and maternal brood care across Arachnopulmonata. Empty symbols indicate absences. For book lungs, half-filled symbol for Schizomida reflects loss of posterior book lung pair; gradient in Araneae reflects transformation of posterior book lung pair to tracheal tubules in derived spiders. For *promenade á deux*, half-filled symbol for Pseudoscorpiones reflects retention only in Cheliferoidea. (b) A mating pair of *Conicochernes crassus* performing the *promenade á deux* (Denmark, Western Australia; photograph: A.Z. Ontano). Maternal brood care in a chenetid (photograph: G. Giribet). (d) *Promenade á deux* behavior in the buthid scorpion *Babycurus gigas* (photograph: M. Cozijn). (e) Maternal care in the vaejovid scorpion *Vaejovis zapoteca*, with scorplings on the back of the female (photograph: C.E. Santibáñez-López). (f) Postembryos of an undescribed species of the schizomid genus *Rowlandius*; hatchling cluster removed from the female’s back for image clarity (photograph: L. Carvalho). (g) A mating pair of the thelyphonid *Mastigoproctus giganteus* (photograph: A. Hochberg, R. Hochberg). (h) Female of the lycosid spider *Hogna* sp. with spiderlings on the back of the female (photograph: J.A. Ballesteros). (i) Female of the whip spider *Phrynus marginemaculatus* with postembryos on the back of the female; yellow marking is a biological paint used to distinguish individuals in a captive breeding colony (photograph: G. Gainett).

Separately, an arachnopulmonate affinity for pseudoscorpions suggests that both a courtship behavior and a mode of parental care are ancient across this group. Like scorpions, Amblypygi, and Thelyphonida, pseudoscorpions of the superfamily Cheliferoidea perform a characteristic courtship dance (the *promenade à deux*), wherein the male clasps the female using the pedipalps and the pair navigate over a substrate (fig. 9b, 9d, 9g; Gravely 1915). The inferred purpose of this behavior is to guide the female to the spermatophore deposited by the male onto the substrate. The *promenade à deux* behavior is secondarily lost in spiders, which exhibit other, often complex, courtship behaviors. In addition, most spiders do not produce an external spermatophore during mating; typically, sperm are passed to specialized copulatory bulbs on the distal palps, which are used for internal fertilization. Given the tree topology supported by analyses (reciprocally monophyletic Panscorpiones + Tetrapulmonata), and under accelerated transformation of character states (shown in fig. 9), the *promenade à deux* appears to be a synapomorphy of Arachnopulmonata that was secondarily lost in spiders as well as in the common ancestor of Pseudoscorpiones, with a secondary regain in Cheliferoidea, or its retention in Cheliferoidea represents a plesiomorphy that reflects arachnopulmonate affinity. An equally parsimonious scenario (under delayed transformation; not shown) constitutes independent gains in Pedipalpi and Panscorpiones, with the same sequence of loss and regain of this character within Pseudoscorpiones. A less ambiguous reconstruction is the presence of a stalked spermatophore attached to the substrate is found across pseudoscorpion superfamilies, as well as in scorpions, Amblypygi, Thelyphonida, and Schizomida (Shultz 2007). A similarity of spermatophore structure in scorpions and pseudoscorpions has previously been noted as well (Francke 1979).

Many pseudoscorpion superfamilies will produce a brood sac on the underside of the female’s opisthosoma that is secreted by genital glands, wherein embryos develop until hatching (fig. 9c). A condition unique to brooding pseudoscorpion lineages is that developing embryos are additionally provisioned by nutritive secretions of the female (Weygoldt 1969). The production of a brood sac from genitalic glands is shared by Amblypygi, Thelyphonida, and Schizomida, which also brood embryos on the underside of the opisthosoma (Gravely 1915; Rowland 1972). The incidence of this mode of development in pseudoscorpions was previously thought to represent a morphological convergence (Shultz 1990). Scorpions exhibit a derived state in this regard, with all extant Scorpiones bearing live young (fig. 9d). Upon birth or hatching from the egg, postembryos of scorpions, Amblypygi, Thelyphonida, and Schizomida will climb onto the female’s back until they advance to additional instar stages (fig. 9f, 9h, 9i). Pseudoscorpion postembryonic care is variable across this order, but can take the form of females forming brood chambers and cohabiting these with offspring (Weygoldt 1969). As with insemination, spiders again bear a derived form of brood care within arachnopulmonates, with the female typically enveloping egg masses in silk. Brood care in spiders is variable; egg sacs may be guarded by females in burrows until juveniles achieve a later instar disperse (e.g., mesotheles; mygalomorphs), attached to webs (most araneomorphs), or carried on the female’s back (e.g., Lycosoidea; fig. 9h).

Given the distribution of the *promenade à deux*, the stalked spermatophore, the production of the maternal brood sac from genitalic glands, and comparable forms of maternal brood care across Chelicerata, we infer these four characters to be ancestral to Arachnopulmonata. As the the oldest known arachnopulmonate, *Parioscorpio venator*, is Silurian in age (439 Mya; Wendruff et al. 2020), the *promenade à deux* may constitute the oldest known courtship behavior.

The recovery of Panscorpiones precipitates reevaluation of other characters, whose homology in now in question. Key among these are venoms of Iocheirata (a clade of venomous pseudoscorpions, which excludes Chthonioidea and Feaelloidea), scorpions, and spiders. As the venom glands of each of these groups do not share positional homology (pedipalpal fingers in pseudoscorpions; posterior-most somite in scorpions; chelicerae in spiders), it is most likely that each group has undergone independent recruitment of housekeeping genes to serve as venom peptides, though striking similarities exist in some toxins of these three groups and may constitute a deep homology (Santibáñez-López et al. 2018; Krämer et al. 2019). On the other hand, the evolution of silks, which occur in spiders, some pseudoscorpions, and some Acariformes (once again, with no shared positional homology of silk-producing organs), is most likely to reflect independent evolutionary gains.

### Prospects for a resolved “arachnid” phylogeny

Topological uncertainty in chelicerate phylogeny extends to the traditionally accepted monophyly of Arachnida, with an array of phylogenomic analyses recovering the derived placement of Xiphosura as the sister group of Ricinulei (Ballesteros and Sharma 2019; Ballesteros et al. 2019). This result has been challenged by another suite of phylogenomic studies (Lozano et al. 2019; Howard et al. 2020) that have suggested three potential solutions to recovering arachnid monophyly: denser taxonomic sampling (Lozano et al. 2019), the use of the site heterogeneous CAT model (Lozano et al. 2019; Howard et al. 2020), and the use of slowly evolving (and/or less saturated) loci (Lozano et al. 2019; Howard et al. 2020). Given the unstable support for arachnid monophyly across phylogenomic data sets, it has been contended that the morphological result of arachnid monophyly should be accepted as the most likely evolutionary scenario (Howard et al. 2020).

As we have previously shown, the most taxon-rich phylogenomic dataset of chelicerates— and the sole analysis sampling all extant chelicerate orders—does not support arachnid monophyly, including under the CAT model (Ballesteros et al. 2019). Recent reanalyses of datasets that had previously recovered arachnid monophyly under certain models (e.g., Regier et al. 2010; 500-slowest evolving genes in Sharma et al. 2014), showed that higher support for Arachnida could be obtained if these were analyzed under a CAT + Poisson model (Howard et al. 2020). But the choice of the model in those reanalyses is peculiar, given that CAT + Poisson has been shown to be demonstrably less accurate than CAT + GTR + Γ_4_ (Halanych et al. 2017). In addition, analyses computed under the CAT + GTR + Γ_4_ model do not consistently recover arachnid monophyly either, including for datasets restricted to slowly-evolving genes (figure 7 of Sharma et al. 2014; Ballesteros and Sharma 2019; Ballesteros et al. 2019).

Across Chelicerata, a subset of genes supporting arachnid monophyly, as identified by a ΔGLS framework, were previously shown to be statistically indistinguishable from the majority (which supported Xiphosura as derived), with respect to 70 parameters, including evolutionary rate, compositional heterogeneity, and alignment length (figure 3 of Ballesteros and Sharma 2019). In the present study, of the 91 phylogenetic analyses we performed using an independent orthology criterion for locus selection (BUSCO genes), not one analysis recovered arachnid monophyly. In addition, surveys of miRNAs revealed no support for Arachnida, either in the form of miRNAs unique to arachnids, or evidence of an arachnid-specific duplication (note that while not all chelicerate orders are represented by genomes, this should not hinder the recovery of putative arachnid-specific miRNAs in our analysis; Garb et al. 2018). Recovering arachnid monophyly in molecular datasets appears to require a concerted, and perhaps contrived, effort to circumscribe taxa, loci, models, and algorithms that will recover this preferred relationship. As we have previously shown, this practice is questionable because it can be used to justify nonsensical groupings (fig. 8 of Ballesteros and Sharma 2019). The attribution of arachnid non-monophyly to unspecified systematic biases or artifacts remains an unsubstantiated notion.

Strong arguments in favor of arachnid monophyly remain the domain of morphological and paleontological datasets; these span the nature of mouthparts, eyes, respiratory systems, and stratigraphic distributions of marine versus terrestrial lineages, among others (reviewed by Howard et al. 2020). Such discussions eerily echo arguments once advanced in support of Tracheata (Myriapoda + Hexapoda, or the terrestrial mandibulates), a group revealed to by molecular phylogenetics to be an artifact of morphological convergence in another subset of terrestrial arthropods. As the history of hypotheses like Pulmonata (Gastropoda) and Tracheata has repeatedly shown, terrestrial lineages are highly prone to convergence, often to an astonishing degree (Friedrich and Tautz 1995; Shultz and Regier 2000; Giribet et al. 2001; Jörger et al. 2010). Shared reduction of the appendage-less intercalary segment (third head segment), the incidence of uniramous appendages, and the organization of the tracheal tubules in hexapods and myriapods serve as powerful examples of how parallel adaptations to life on land can confound interpretations of synapomorphies.

We submit that an objective approach to testing phylogenetic hypotheses of terrestrialization in arthropods must regard traditional groupings with skepticism, rather than querying molecular sequence data for genes and datasets supporting preconceived relationships. Such investigations must also account for new neurophylogenetic characters that have recently suggested morphological support for a closer relationship of Xiphosura to Arachnopulmonata (Lehmann and Melzer 2019a, 2019b). Due to the lack of genomes for Ricinulei (the putative Xiphosura sister group in some phylogenies) as well as other poorly studied arachnid groups (e.g., Palpigradi, Solifugae), we were not able to assess miRNAs or other rare genomic changes to test the competing hypothesis of Ricinulei + Xiphosura. However, the incidence of whole genome duplications in horseshoe crabs proffers the tantalizing possibility of applying the approaches used herein to assess this competing hypothesis, as at least one of the two WGD events in Xiphosura is thought to be ancient. The discovery of shared duplications of gene families, miRNAs, and syntenic blocks between different sets of chelicerate orders could be used to evaluate independently the monophyly of Arachnida, as well as the placement of the unstable apulmonate orders. Future efforts should therefore target the generation of genomic resources for Ricinulei, Palpigradi, and Solifugae to reevaluate such hypotheses as Haplocnemata (Solifugae + Pseudoscorpiones), Megoperculata (Palpigradi + Tetrapulmonata), and Arachnida itself.

## Conclusions

Consilience in phylogenetics is the outcome of multiple, independent topological tests recovering support for the same hypothesis (e.g., Rota-Stabelli et al. 2011; Fröbius and Funch 2018; Marlétaz et al. 2019). Here, we demonstrated that analyses of sequence data, gene family duplications, gene tree topologies of arachnopulmonate-specific paralogs, and miRNA duplications independently support a nested placement of pseudoscorpions within Arachnopulmonata. Our results reinforce that topological accuracy in the placement of long branch taxa is most affected by dense sampling of basally branching lineages, rather than algorithmic approach (supermatrix versus coalescent-based summary methods), matrix completeness, or model choice alone. Improvements to chelicerate phylogeny must therefore focus on the identification of basally branching groups within orders whose internal relationships remain poorly understood, such as Solifugae, Amblypygi, and Uropygi (Thelyphonida + Schizomida). Leveraging rare genomic changes stemming from the genome duplications exhibited by a subset of chelicerate orders may be key to resolving some of the most obdurate nodes in the chelicerate tree of life.

## Materials and Methods

### Species sampling

For phylogenetic reconstruction, we generated a dataset of 117 chelicerates (40 pseudoscorpions, 12 scorpions, 17 spiders, 4 Pedipalpi, 13 Opiliones, 5 Ricinulei, 3 Xiphosura, 2 Solifugae, 9 Parasitiformes, 10 Acariformes, 2 Pycnogonida) and 15 outgroups (3 Onychophora, 4 Myriapoda, 8 Pancrustacea), with most of these taxa sequenced previously by us. Taxon selection prioritized the representation of basal splits in all major groups (Sharma et al. 2015c; Fernández et al. 2017, 2018; Ballesteros et al. 2019, 2020; Santibáñez-López et al. 2019, 2020; Benavides et al. 2019). Libraries of high quality were additionally selected such that all chelicerate orders were represented in >95% of loci in all matrices constructed. While we trialed the inclusion of a palpigrade library recently generated by us (Ballesteros et al. 2019), the low representation of BUSCO genes for this taxon across datasets (46-70%) prohibited the inclusion of this order in downstream analyses. A list of taxa and sequence accession data is provided in table S1.

### Orthology inference and phylogenomic methods

Candidate ORFs were identified in transcripts using TransDecoder (Haas et al. 2013). Loci selected for phylogenomic analysis consisted of the subset of 1066 Benchmarked Universal Single Copy Orthologs identified for Arthropoda (BUSCO-Ar). For each library, these were discovered using a hidden Markov model approach, following the procedure detailed in Leite et al. (2018). Multiple sequence alignment was performed using MAFFT 7.3.8 *(– anysymbol –auto*; Katoh and Standley 2013). Gap-rich regions were masked with trimAl 1.2 *(–gappyout*; Capella-Gutiérrez et al. 2009) and alignment coverage verified and sanitized with Al2Phylo *(-m 50 -p 0.25 -t 20*; Ballesteros and Hormiga 2016).

To assess the tradeoff between data completeness and number of loci per dataset, six matrices were constructed by setting taxon occupancy thresholds to 55% (1002 loci), 60% (945 loci), 65% (846 loci), 70% (693 loci), 75% (480 loci), and 80% (248 loci) of total taxa. These thresholds were selected to represent broadly commonly occurring values for matrix completeness in phylotranscriptomic studies of metazoans. Representation of each terminal and ordinal lineage per matrix is provided in table S2.

To assess the effect of denser taxonomic sampling on the placement of Pseudoscorpiones, basally branching lineages of pseudoscorpions (corresponding to superfamilies or families) were sequentially pruned until only Cheliferoidea (Cheliferidae + Chernetidae) was retained. Thus, six additional matrices were constructed, with sequential pruning of Chthonioidea (6 terminals), Feaelloidea (2 terminals), Neobisioidea (10 terminals), Garypoidea (5 terminals), Garypinoidea (3 terminals), and Cheridoidea + Sternophoroidea (two terminals). Pruning was performed for each of the six matrices constructed according to taxon occupancy thresholds, resulting in 42 matrices in total.

Tree topologies for individual loci and for concatenated datasets were computed with IQ-TREE 1.6.8 (Nguyen et al. 2015; Chernomor et al. 2016), coupled with model selection of substitution and rate heterogeneity based on the Bayesian Information Criterion (Kalyaanamoorthy et al. 2017) and 1000 ultrafast bootstraps to assess branch support *(-m MFP -mset LG, JTT, WAG -st AA -bb 1000*; Hoang et al. 2018). For the subset of least complete matrices (55% taxon occupancy), we additionally performed model selection under the posterior mean site frequency (PMSF) (LG + C20 + F + Γ), a mixture model that approximates the CAT model in a maximum likelihood framework (Lartillot and Philippe 2004; Wang et al. 2018).

For phylogenetic analyses using multispecies coalescent methods, species trees were estimated with ASTRAL v. 3 (Mirarab and Warnow 2015; Zhang et al. 2018), using gene trees from IQ-TREE analyses as inputs. Phylogenetic signal at the level of individual genes was quantified using the gene-wise log-likelihood score (ΔGLS) for the unconstrained tree versus a competing hypothesis (Pseudoscorpiones + Acariformes; Pseudoscorpiones + Parasitiformes; Pseudoscorpiones + Scorpiones) (Shen et al. 2017). This metric maps the relative support for each of two competing hypotheses, for every locus in the dataset; the amplitude of the log-likelihood indicates the degree of support for either hypothesis.

### Embryo collection, sequencing, and mapping of homeodomains

Given that transcriptomes of adult tissues have been shown to sample poorly transcription factors relevant for developmental patterning in arachnids (Sharma et al. 2016), assessment of homeodomain duplications was performed only for genomes and developmental transcriptomes. The genome of *C. scorpioides* was excluded from this analysis, due to the fragmentation of the assembly.

*C. crassus* (Pseudoscorpiones: Chernetidae) were hand collected from underneath the bark of karri trees in Denmark, Western Australia (−34.963640, 117.359720). Individuals were reared in plastic containers containing damp paper towels at room temperate to simulate living conditions between bark and sapwood. Adult pseudoscorpions were fed a combination of cricket nymphs and *ap*^*−/−*^ fruit flies. Females of *C. crassus* carry developing embryos in a brood sac on the underside of the opisthosoma; individuals were checked for the presence of embryos. Females carrying embryos were separated from the colony for 12-72 hours to prevent cannibalism and allow embryos to mature to a range of developmental time points. Entire brood sacs were then separated from the opisthosoma using forceps wetted with distilled water to prevent damage to the females before being returned to the colony.

Establishment of *Phrynus marginemaculatus* (Amblypygi: Phrynidae) for study of developmental genetics and comparative development was previously described by Gainett and Sharma (2020). Embryos of the whip spiders *Charinus ioanniticus* and *Charinus israelensis* were obtained by hand collecting brooding females from two cave sites in Israel, Hribet Hruba (31.913280, 34.960830) and Mimlach (32.858150, 35.44410). Two stages of deutembryos were obtained and sequenced for each species. Further details are provided in Gainett et al. (2020).

Field collection of embryos of the tarantula *Aphonopelma hentzi* (Araneae: Theraphosidae) for developmental genetics and transcriptomics was previously described by Setton et al. (2019).

Field collection of embryos and larvae was performed for five species of Pycnogonida: *Nymphon moelleri* (Nymphonidae), *Pallenella flava* (Callipallenidae), Stylopallene cheilorhynchus (Callipallenidae), *Phoxichilidium femoratum* (Phoxichilidiidae), and *Tanystylum orbiculare* (Ammotheidae). Details of collection and sequencing are provided in Ballesteros et al. (2020).

Embryos were transferred to Trizol Tri-reagent (Ambion Life Technologies, Waltham, MA, USA) for RNA extraction, following manufacturer’s protocols. Library preparation and stranded mRNA sequencing were performed at the University of Wisconsin-Madison Biotechnology Center on an Illumina HiSeq 2500 platform (paired-end reads of 125 bp). Raw sequence reads are deposited in NCBI Sequence Read Archive. Filtering of raw reads and strand-specific assembly using Trinity v. 2.8.3 followed our previous approaches (Sharma et al. 2014; Ballesteros et al. 2019).

Discovery of homeobox genes followed the approach previously outlined by Leite et al. (2018). Briefly, homeodomain sequences were identified from genomes and embryonic transcriptomes using BLAST v. 2.9.0 or v. 2.10.0 (tblastn) (Altschul et al. 1990). Queries consisted of amino acid homeodomain sequences from outgroup arthropod species in HomeoDB (Zhong et al. 2008; Zhong and Holland 2011) combined with homeodomain sequences from *Parasteatoda tepidariorum* (Schwager et al. 2017), *Centruroides sculpturatus* (Schwager et al. 2017), *Mesobuthus martensii* (Cao et al. 2013), and *Strigamia maritima* (Chipman et al. 2014). As additional chelicerate ingroup taxa, we included the genome of the horseshoe crabs *Limulus polyphemus* (Kenny et al. 2016) and *Carcinoscorpius rotundicauda* (Shingate et al. 2020), the genomes of the mites *Tetranychus urticae* (Grbic et al. 2011) and *Galendromus occidentalis* (Hoy et al. 2016), and a recently re-sequenced embryonic transcriptome of the harvestman *Phalangium opilio* (SRX450969; Sharma et al. 2012; Ballesteros and Sharma 2019). As additional outgroup taxa, we included the embryonic transcriptomes of the millipede *Glomeris marginata* and the onychophoran *Euperipatoides kanangrensis* (Janssen and Budd 2013). We thus assessed homeobox gene duplication for 26 panarthropod species.

All initial BLAST hits were retained. Next, the full protein sequences of the BLAST hits were predicted with TransDecoder v. 5.5.0 (Haas et al., 2013) with default parameters (*-m 100*; predicted transcripts with less than 100 amino acids were not retained), and thereafter analyzed using the Conserved Domain Database (CDD) (Marchler-Bauer et al. 2015) to confirm the presence of homeodomains and annotate other functional domains. BLAST hits that did not have homeodomains identified by CDD were removed. Transcripts within a species that had identical protein sequences predicted to encode homeodomains were manually checked. Because this approach conservatively emphasized retention of complete homeobox genes with conserved sequences, we cannot rule out the exclusion of partial transcripts of homeobox genes that lack homeodomains or orthologs with highly divergent sequences. Multiple sequence alignment, trimming to retain only the homeodomain, and classification of verified homologs followed procedures described by Leite et al. (2018).

### Analysis of appendage patterning ohnologs

Homologs of four appendage patterning genes were retrieved from the *C. crassus* transcriptome using approaches described above. Multiple sequence alignment of peptide sequences and alignment trimming followed the approach of Nolan et al. (2020). Maximum likelihood inference of tree topologies was performed using IQ-TREE under an LG + I + Γ substitution model. Nodal support was estimated using ultrafast bootstrapping.

### *Cordylochernes scorpioides* genome sequencing

Illumina fragment libraries (insert sizes 270b and 420b) and mate-pair libraries (insert sizes 2kb, 4kb and 8kb) were constructed by Lucigen Corporation (Middleton, WI, USA). Fragment libraries were constructed from genomic DNA extracted from single individual inbred males; to meet DNA input requirements for mate-pair library construction, genomic DNA from 12 4^th^-generation inbred individuals was pooled. Fragment libraries were sequenced on HiSeq X with 150b paired-end sequencing (Hudson Alpha Genomic Services Lab, Huntsville AL), and mate-pair libraries were sequenced on MiSeq with 150b paired-end sequencing at Lucigen Corporation. The read data was assembled *de novo* at 125X coverage using MaSuRCA v. 3.2.3 (Zimin et. al. 2013), with additional scaffolding using SSPACE Standard v 3.0 (BaseClear BV, Netherlands) followed by gap-filling using GapFiller v1.12 (BaseClear). The draft *C. scorpioides* genome assembly was submitted to GenBank (GenBank: QEEW00000000.1) and read data were deposited in NCBI SRA (SRA:SRP144365 BioProject:PRJNA449764).

### microRNA and Hox genes orthology search

Previous work on miRNA occurrence in the genome of the house spider *Parasteatoda tepidariorum* identified 40 miRNA families shared across Arthropoda, and a further 31 either unique to spiders (n=30) or unique to arachnopulmonates (n=1) (Leite et al. 2016). To extend this survey to new taxa, we searched for miRNA families in the draft genome assembly of *C. scorpioides* (GCA_003123905.1), as well as the genome of *Mesobuthus martensii* (GCA_000484575.1). All miRNA reported from *P. tepidariorum* were retrieved from the miRBASE and used as query sequences (Kozomara et al. 2019). An initial BLAST search was performed (*blastn –word_size 4 –reward 2 –penalty –3 –evalue 0.05*) and sequences with e-value < 0.05 and percentage identity > 70% were retained. To accommodate the fragmentation of the *C. scorpioides* genome, as well as heterozygosity, putative hits were retained only if both the ELEKEF and KIWFQN motifs were discovered in the peptide translation, and peptide sequences were unique (i.e., pairs of sequences with only synonymous substitutions were considered putative alleles). Putative homologs were verified by multiple sequence alignment using MAFFT v. 7.407 (Katoh and Standley, 2013). The structure and minimum free energy of these selected miRNAs were analyzed with RNAfold v. 2.4.13 (as part of the ViennaRNA Package 2.0; Lorenz et al. 2011) and with The Vienna RNA WebServer (http://rna.tbi.univie.ac.at/cgi-bin/RNAWebSuite/RNAfold.cgi) using default settings. Regarding the previous survey of miRNA families in 16 ecdysozoan taxa by Leite et al. (2016), we corroborated all reported results, except for the discovery that the mygalomorph spider *A. hentzi* exhibits only a single copy of the miRNA *pte-bantam*.

## Acknowledgements

We are indebted to Leonardo Carvalho, Adele Hochberg, Rick Hochberg, and Gonzalo Giribet for contributing photographs of Schizomida, Thelyphonida, and Pseudoscorpiones. Sequencing of *C. crassus* was performed at BioTechnology Center at UW-Madison. Access to computing nodes for intensive tasks was provided by the Center for High Throughput Computing (CHTC) and the Bioinformatics Resource Center (BRC) of the University of Wisconsin–Madison. We thank La Autoridad Nacional del Ambiente for permission to collect *C. scorpioides* in the República de Panamá and the Smithsonian Tropical Research Institute for logistical support.

## Author contributions

A.Z.O. and P.P.S. conceived of the study. A.Z.O., P.P.S., and M.S.H. collected specimens of pseudoscorpions in the field. M.S.H. performed taxonomic identification. A.Z.O. cultivated pseudoscorpion embryos, performed the sequencing, and implemented phylogenomic analyses. Fieldwork and tissue collection was performed by S.A., J.AB., G.G., E.G.R., and P.P.S. for whip spider embryos; by G.B. and P.P.S. for sea spider embryos; and by E.V.W.S. and P.P.S. for mygalomorph embryos. Analysis of homeobox duplications was performed by A.Z.O., G.G., K.F.C., J.A.B., E.V.W.S., J.T.Z., and P.P.S. Analysis of miRNAs was performed by C.E.S.L. J.A.Z. and D.W.Z. collected *C. scorpioides* in Panamá, established a laboratory population of the pseudoscorpion, and conceived of the genome sequencing of *C. scorpioides*. S.M., J.A.Z. and D.W.Z. were responsible for *C. scorpioides* genome sequencing and assembly. A.Z.O. and P.P.S. wrote the manuscript, and all authors edited and approved the final content.

## Data availability

The complete dataset, including sequence alignments, tree files, miRNA alignments, and embryonic transcriptomic assemblies, have been deposited on the Dryad Digital Repository. Raw read data are available in NCBI Sequence Read Archive. The *C. scorpioides* genome assembly and associated SRA are available in GenBank under the WGS master record QEEW00000000.1.

## Permitting

Specimens of *C. crassus* were collected in Western Australia under permit number 08‒000214‒6 from the Department of Parks and Wildlife. Specimens of *C. scorpioides* were collected in Panamá under permits SE/A-92-05 (collecting) and SEX/A-142-05 (export), from the Autoridad Nacional del Ambiente, Repú blica de Pamaná; and permit number 68818 (quarantine) from the Ministerio de Desarrollo Agropecuario, República de Panamá.

---

**Figure S1.**
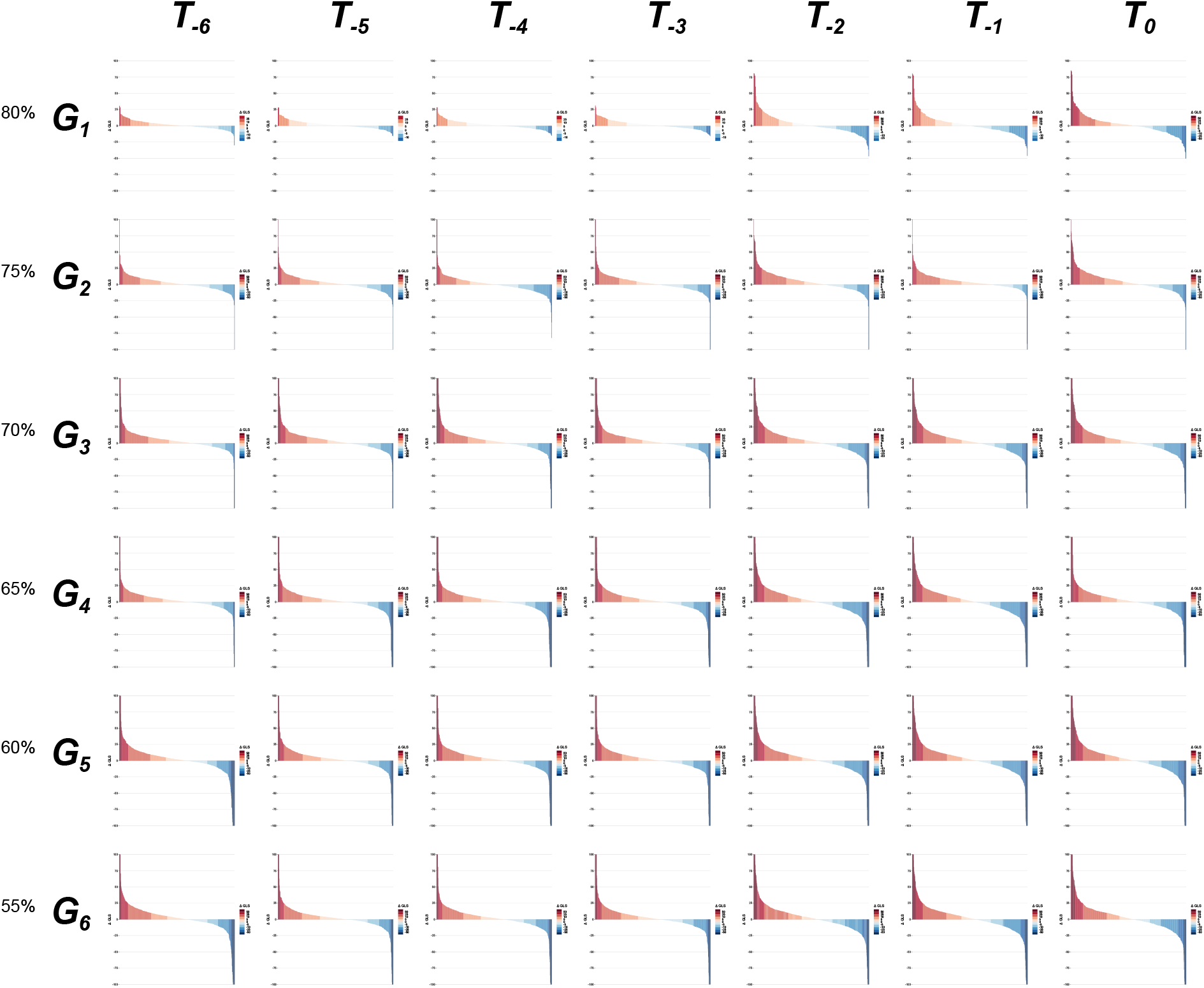
Distribution of ΔGLS scores and amplitudes across 42 datasets.

**Table S1.** Study taxa and accession data.

**Table S2.** Representation of species and orders per matrix (in number of loci).

**Table S3.** Tabulation of homeobox genes discovered across 26 Panarthropoda.

